# Discovering the Molecular Determinants of *Phaeobacter inhibens* susceptibility to *Phaeobacter* phage MD18

**DOI:** 10.1101/2020.04.13.038638

**Authors:** Guillaume Urtecho, Danielle E. Campbell, David M. Hershey, Rachel J. Whitaker, George A. O’Toole

## Abstract

Bacteriophage technologies have immense potential as antibiotic therapies and in genetic engineering. Understanding the mechanisms that bacteriophages implement to infect their hosts will allow researchers to manipulate these systems and adapt them to specific bacterial targets. Here, we isolated a bacteriophage capable of infecting the marine alphaproteobacterium *Phaeobacter inhibens* and dissected its mechanism of infection. *Phaeobacter* phage MD18, a novel species of bacteriophage isolated in Woods Hole, MA, exhibits potent lytic ability against *P. inhibens* and appears to be of the *Siphoviridae* morphotype. Consistent with this finding, the sequence of the MD18 revealed significant similarity to another siphophage, the recently discovered *Roseobacter* phage DSS3P8. We incubated MD18 with a library of barcoded *P. inhibens* transposon insertion mutants and identified 22 genes that appear to be required for phage predation of this host. Network analysis of these genes using genomic position, GO term enrichment, and protein associations reveals that these genes are enriched for roles in assembly of a type IV pilus (T4P) and regulators of cellular morphology. Our results suggest that T4P serve as receptors for a novel marine virus that targets *P. inhibens*.

**Importance:** Bacteriophages are useful non-antibiotic therapeutics for bacterial infections as well as threats to industries utilizing bacterial agents. This study identifies *Phaeobacter* phage MD18, the first documented phage of *Phaeobacter inhibens*, a bacterium with promising use as a probiotic for aquatic farming industries. Genomic analysis suggests that the *P*

*haeobacter* phage MD18 has evolved to enhance its replication in *P. inhibens* by adopting favorable tRNA genes as well as through genomic sequence adaptation to resemble host codon usage. Lastly, a high-throughput analysis of *P. inhibens* transposon insertion mutants identifies genes that modulate host susceptibility to phage MD18 and implicates the type IV pilus as the likely receptor recognized for adsorption. This study marks the first characterization of the relationship between *P. inhibens* and an environmentally sampled phage, which informs our understanding of natural threats to the bacterium and may promote the development of novel phage technologies for genetic manipulation of this host.

## Introduction

Viruses are the largest source of genetic diversity known. As such, they have given rise to an incredible assortment of potential therapeutic and genetic tools used as sustainable substitutes for antibiotics(1, 2), targeted bacterial delivery systems(3), and for bacterial engineering(4). However, the unique ecological relationships and infection mechanisms that have evolved in the wake of the phage-host arms race are diverse and complex, making it difficult to systematically leverage these systems for practical purposes. Further studies of phage-host interactions are essential to understand and manipulate the mechanisms phages implement to infect their host.

*Phaeobacter inhibens* is a marine bacterium and member of the *Rhodobacteriales* order of Alphaproteobacteria. Organisms in this clade are found worldwide, especially near coastal waters(5, 6), and produce a diverse array of secondary metabolites(7, 8). *P. inhibens* is most known for the production of the antibiotic tropodithietic acid (TDA), which protects its natural algal symbiont from marine pathogens(8, 9). Due to this property, *P. inhibens* serves as a useful probiotic in oyster and other aquatic farms to prevent colonization by pathogenic *Vibrio* species(10, 11). Despite its importance in the aquatic farming industry, viral predators that target this microbe remain poorly understood. Characterizing phage-host interactions within this industrially relevant host will enable researchers to implement phage engineering approaches for this species and provide insight into potential natural threats of *P. inhibens*.

Here, we first isolated a siphophage capable of infecting *P. inhibens* from an aquatic environment in Woods Hole, MA and characterized this phage using transmission electron microscopy (TEM) and whole-genome sequencing. Based on the morphology and genome sequence, this isolate represents a novel siphophage, which we name *Phaeobacter* phage MD18. We used a barcoded transposon insertion mutant library (i.e., BarSeq) to characterize the relationship between *Phaeobacter* phage MD18 and *P. inhibens*. The type IV pilus system, the ChvI/ChvG two-component system, and regulators of cell division were identified as key determinants of infection. This work characterizes the genetic basis of phage infection in an underexplored non-model host system.

## Results

### Isolation and characterization of *Phaeobacter inhibens* phage MD18

To isolate a natural bacteriophage capable of infecting *P. inhibens*, we obtained samples from twenty aquatic environments around Woods Hole, MA. We filtered these samples to isolate phage particles and applied the resulting filtrates to exponentially growing liquid cultures of *P. inhibens*. To enrich for phages infecting *P. inhibens*, we applied the resulting filtrates to exponentially growing liquid cultures of *P. inhibens* and incubated the bacteria-phage mixtures overnight. After filtering enrichments as before, we spotted each lysate onto a lawn of *P. inhibens* grown on agar plates and monitored for the formation of plaques. Of the 20 enriched phage samples, one sample sourced from a seashore environment produced a clear plaque, suggesting the presence of a lytic phage. This enriched phage sample contained high titers of plaque forming units (∼10^10^ PFU/mL; **Figure 1A**) and repressed host growth in liquid culture at concentrations of the phage below the limit of detection by plaquing (<200 PFU/ml; **Figure 1B**). Growth inhibition occurred after incubation of the phage lysate with chloroform, confirming that predatory cellular microbes (*e*.*g. Bdellovibrio* spp.) were not responsible for this phenomenon. Propagation of clear plaques indicated essentially complete bacterial host lysis, and thus was likely the result of a lytic phage, henceforth referred to as *Phaeobacter* phage MD18.

**Figure 1.**
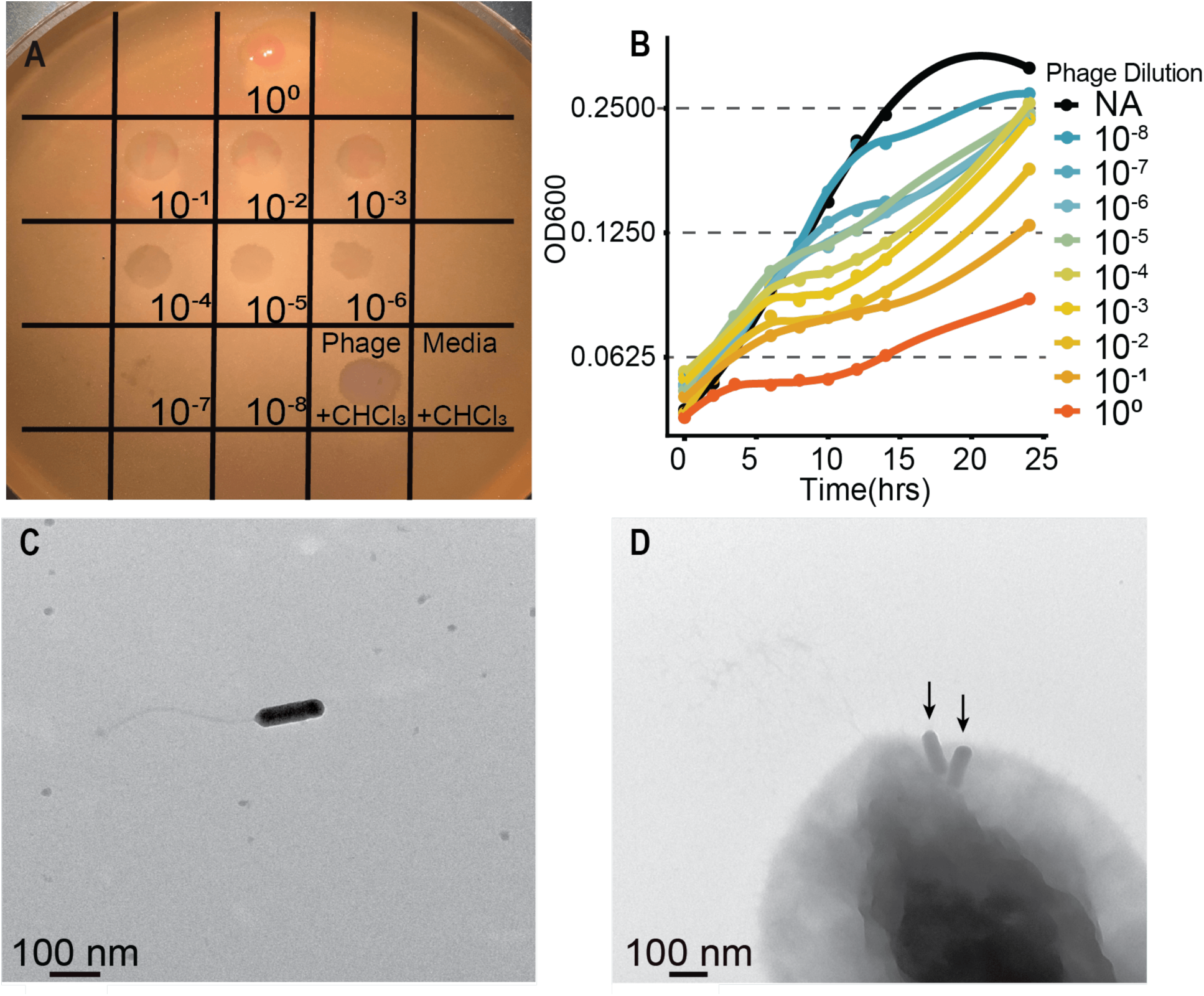
Characterization of Phage MD18. **A)** Host lysis after top agar spotting of *Phaeobacter* phage MD18 onto lawn of *P. inhibens*. Approximately 5 plaques are detected in the 10^−7^ dilution. Given that 5 μl of the lysate was spotted here, this corresponds to a phage titer of ∼10-10 PFU/ml. Chloroform-treated control (+CHCl_3_), spotted here as the undiluted lysate, suggests that cellular predators are not responsible for plaque formation. Media control also underwent chloroform treatment. **B)** Growth of *P. inhibens* in the presence of the indicated titer of phage MD18, as monitored by optical density at 600 nm (OD_600_). NA, none added, is the no phage control. Each line represents the average of three growth curves performed in a 96 well plate. See Material and Methods for additional details. **C)** Transmission electron microscopy (TEM) identifies phage particles resembling *Siphoviridae*. **D)** TEM image of phage MD18 (arrows) adsorbing to the surface *P. inhibens*. Adsorbed phage particles are adjacent to apparent pilus-like structures. A scale bar is shown in panels C and D.

To analyze phage MD18 morphology, we visualized filter-sterilized phage lysates by TEM. TEM revealed phage particles with prolate heads roughly 100 nm in length, within a normal range for bacteriophages(12) (**Figure 1C**). The long, flexible, non-contractile tail suggests that phage MD18 belongs to the *Siphoviridae* morphotype(13). Furthermore, we observed incidences of MD18 particles adsorbed to the surface of *P. inhibens* cells, positioned roughly at the bacterial cell pole (**Figure 1D**). Together, these results suggest that phage MD18 is a potent lytic phage of the *Siphoviridae* morphotype and may use a structure at the cell pole for host recognition.

### Genome Sequence of *Phaeobacter* phage MD18 and evidence of host adaptation

We further characterized phage MD18 by performing whole-genome shotgun sequencing. We assembled a single circular contig, which was 149,262 bp in length. A BLAST search of this assembly did not reveal extensive similarity with published viral genomes. However, the best match corresponded to a recently discovered *Roseobacter* phage DSS3P8 (KT870145.1) which is also of the *Siphoviridae* morphotype(14) (**Figure S1**). In light of the classification guidelines established by the International Committee on Viral Taxonomy’s Bacterial and Archaeal Viruses Subcommittee, *Phaeobacter* phage MD18 represents a novel genus, we name ‘*Kasiahvirus*’ within the family of *Siphoviridae(15)*. We then annotated this genome using RAST(16, 17) and identified 257 putative genes (**Supplementary Table S1**), of which 215 represented ‘hypothetical proteins’. Notable identified genes include a RecD-like DNA helicase and a DNA polymerase III ε subunit, which are likely components of the phage replication system, as well as a N-acetylmuramoyl-L-alanine amidase, which have been demonstrated to facilitate bacterial cell wall degradation and cell lysis(18, 19).

We used tRNAscan-SE v. 2.0(20, 21) to identify tRNAs in the *Phaeobacter* phage MD18 genome and found 32 tRNA genes (28 unique) (**Supplementary Table S2**). These tRNA genes corresponded to codons which were significantly enriched in the *Phaeobacter* phage MD18 genome compared to codons without a corresponding tRNA gene (**Figure 2A**) (*p* = 6.4 × 10^−5^, Wilcox rank sum test). Phage genomes frequently encode tRNA genes, which facilitate the translation of phage transcripts in host bacteria(22–24). Another assumption of phage genome evolution is that phage codon usage adapts to resemble that of their bacterial hosts(23, 24). Supporting this model, the relative codon preference of all codons in *Phaeobacter* phage MD18 genes was significantly positively correlated with codon usage in *P. inhibens* (*r* = 0.877, *p* < 2.2 × 10^−16^) and less significantly correlated with a distantly-related non-host bacterium, *Escherichia coli (r =* 0.656, *p* = 4.094 × 10^−9^) (**Figure 2B**).

**Figure 2.**
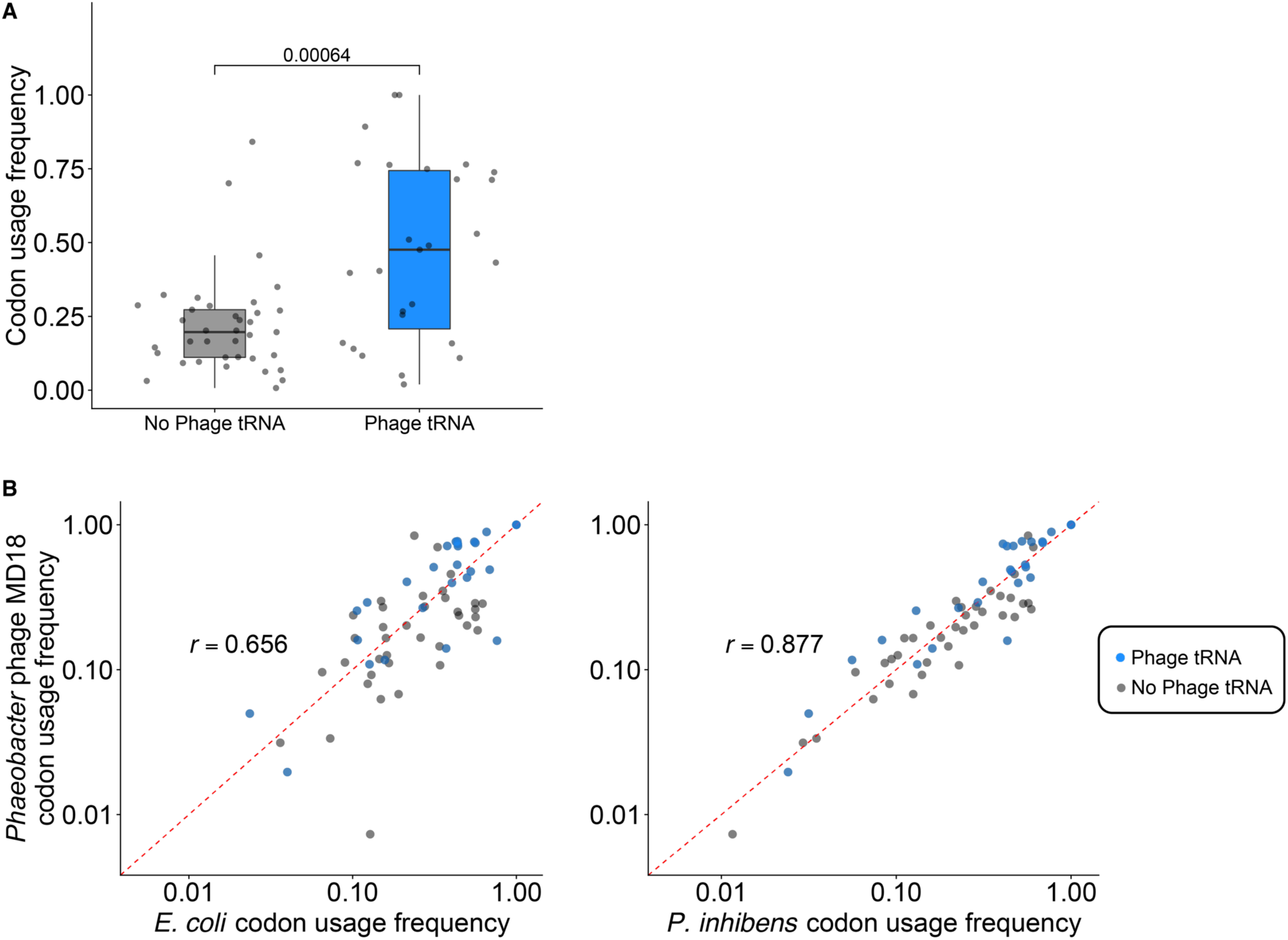
Bacteriophage tRNA genes show adaptation to host and phage genomes. **A)** Codons corresponding to phage-encoded tRNA genes are significantly enriched in the *Phaeobacter* phage MD18 genome compared to codons without a corresponding phage tRNA gene (*p* = 6.4 × 10^−5^, Wilcox rank sum test). **B)** Codon usage frequencies between *Phaeobacter* phage MD18 and *P. inhibens* (right panel, *r* = 0.877, *p* < 2.2 × 10^−16^) exhibit better correlation than *Phaeobacter* phage MD18 to *Escherichia coli* (left panel, *r* = 0.656, *p* = 4.094 × 10^−9^). Blue dots indicate codons with corresponding tRNA genes encoded within the phage genome. Dashed red line illustrates a perfect correlation.

### Identification of MD18 resistant *Phaeobacter inhibens* mutants in a barcoded transposon insertion mutant library

We sought to characterize the relationship between *Phaeobacter* phage MD18 and *P. inhibens* by identifying host gene products that confer susceptibility to infection. Recent work has used transposon insertion sequencing to rapidly perform reverse-genetic screens in hosts and identify genes that contribute to phage susceptibility(25, 26). We selected for *P. inhibens* transposon insertion mutants with decreased susceptibility to phage by exposing a previously constructed, barcoded transposon mutant library(27) to phage MD18 (**Figure 3A**). This library consisted of 205,898 variants, each carrying a randomly barcoded transposon insertion mapped to one of 3,341 *P. inhibens* genes annotated using RAST(16, 17). To impose selection on this library, we incubated the exponentially growing library with *Phaeobacter* phage MD18 isolate (1.5 × 10^8^ PFU) for eight hours, conditions which previously allowed for significantly delayed growth in wildtype *P. inhibens* (**Figure 1B**). In parallel, we grew the same library without phage selection as a comparative control. After growth in triplicate of the phage-exposed and control cultures, we extracted DNA from all samples and sequenced barcoded insertion sites in each population to measure the relative frequency of each *P. inhibens* mutant variant. Concurrently, we plated the post-selection population on agar plates and isolated 12 individual clones. We found that all 12 isolates exhibited a marked increase in resistance to phage challenge (**Figure 3B**).

**Figure 3.**
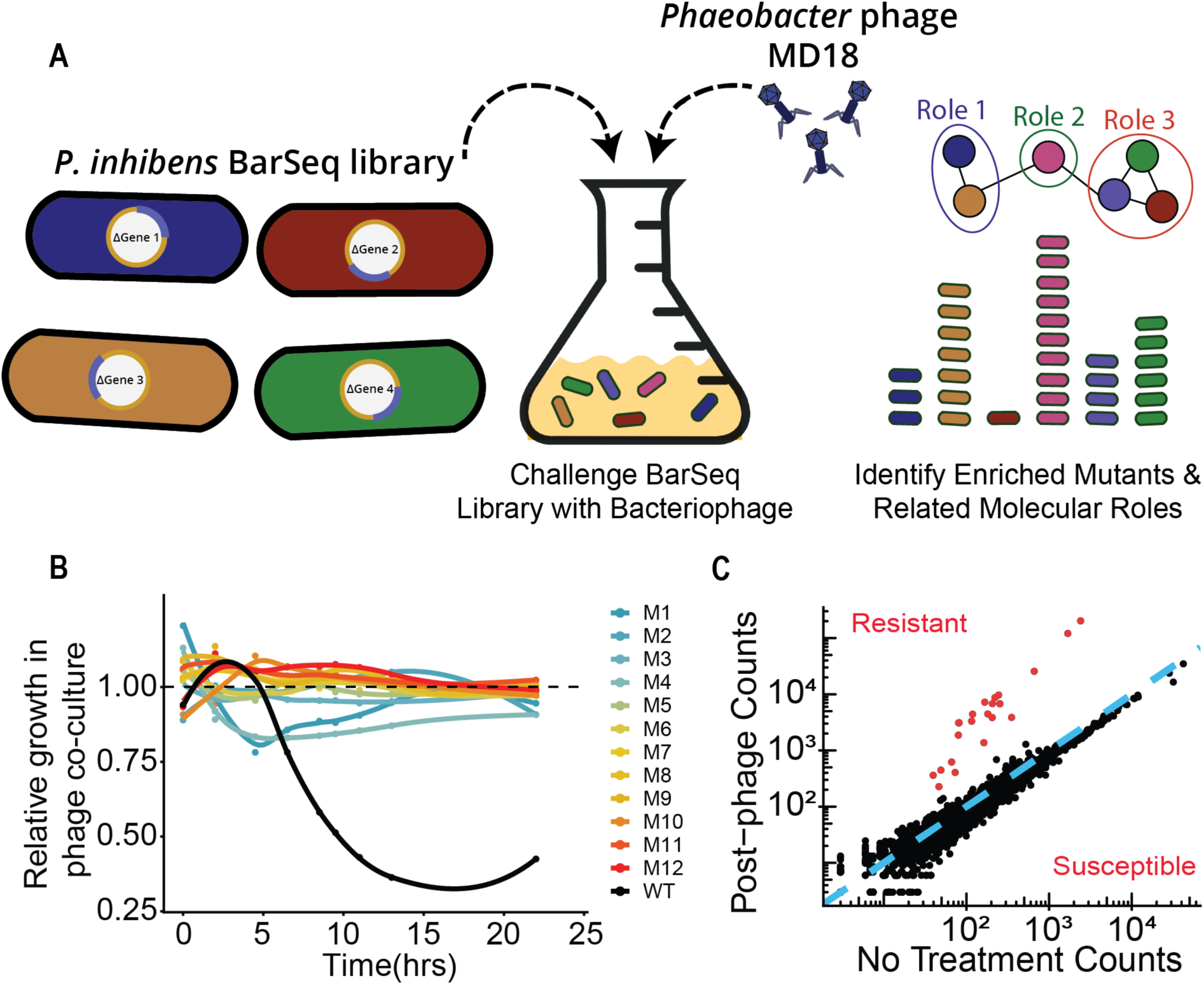
BarSeq screen identifies genetic loci that contribute to phage susceptibility. **A)** Experimental scheme for BarSeq assay. After identification of a phage species capable of infecting *P. inhibens*, this phage was incubated with the BarSeq library and the barcodes of the survivors sequenced. The relative abundance of the barcodes was compared to a parallel culture in which no phage is added, and the functional relationships between those genes analyzed. **B)** Library variants that survived selection (M1-M12; colored lines) are nearly entirely resistant to phage MD18 while WT growth is significantly hindered (black line). Relative growth is the ratio of the grown rate with phage compared to without phage. **C)** Comparison of BarSeq mutant counts grown with and without phage. Mutations in genes above the blue line (y=x) exhibited a relative increase in abundance following phage incubation, indicating enhanced resistance. Highlighted points (red) are considered statistically significant and exhibited fitness scores three standard deviations less than or greater than the mean.

Using the relative abundances of barcodes present in the total cultures with and without phage exposure, we determined the fitness of all mutant strains within the BarSeq library. Of the 3,341 genes for which fitness values were determined, 22 exhibited significantly greater fitness scores when grown in the presence of phage (**Figure 3C, Figure S2**). Interestingly, these genes clustered within four primary operons, suggesting the presence of highly connected functions (**Figure 4A**). GO term analysis using SEED subsystem annotations(17) showed that these significant hits were enriched for genes encoding a type II secretion system involved in pilus formation (**Figure 4B**). Other significant hits included the ChvI/ChvG two-component system and Fts cell division proteins (**Table 1**). We did not identify any genes with negative fitness scores, which would indicate a role in promoting phage resistance.

**Table 1.** Significant genes identified in BarSeq study. Genes are ordered by genomic position and colored according to STRING-assigned cluster. Genomic annotation was performed using RAST with *P. inhibens* DSM 17395. Fitness scores determined using the sum of three biological replicates.

| RAST Annotation | Gene Start | Gene End | Fitness Score |
| --- | --- | --- | --- |
| DNA-binding response regulator ChvI | 108581 | 109282 | 6.18 |
| Sensor histidine kinase ChvG | 109316 | 111028 | 6.39 |
| Peptidase, M23/M37 family | 267155 | 268339 | 3.48 |
| Cell-division-associated, ABC-transporter-like signaling protein FtsE | 337993 | 338670 | 2.97 |
| Cell-division-associated, ABC-transporter-like signaling protein FtsX | 338667 | 339566 | 3.17 |
| Type IV prepilin peptidase TadV/CpaA | 620846 | 621352 | 3.21 |
| Flp pilus assembly protein TadD, contains TPR repeat | 621372 | 622220 | 5.25 |
| Flp pilus assembly protein TadD, contains TPR repeat | 622261 | 622863 | 2.51 |
| Type II/IV secretion system protein TadC, associated with Flp pilus assembly | 622867 | 623847 | 5.06 |
| Flp pilus assembly protein TadB | 623858 | 624826 | 4.81 |
| Type II/IV secretion system ATP hydrolase TadA/VirB11/CpaF, TadA subfamily | 624826 | 626271 | 5.25 |
| Type II/IV secretion system ATPase TadZ/CpaE, associated with Flp pilus assembly | 626301 | 627533 | 4.60 |
| hypothetical protein | 627567 | 627707 | 2.25 |
| Outer membrane lipoprotein precursor, OmpA family | 627734 | 628369 | 4.53 |
| Type II/IV secretion system secretin RcpA/CpaC, associated with Flp pilus assembly | 628380 | 629786 | 5.33 |
| Flp pilus assembly protein RcpC/CpaB | 630014 | 630874 | 5.26 |
| Flp pilus assembly protein, pilin Flp | 631293 | 631478 | 5.20 |
| ABC-type dipeptide transport system, periplasmic component | 896851 | 897450 | 4.77 |
| Flp pilus assembly protein TadG | 897454 | 898014 | 5.51 |
| hypothetical protein | 898011 | 899693 | 5.28 |
| N-acetylmuramoyl-L-alanine amidase | 2107483 | 2108760 | 3.19 |
| CPN protein | 2482842 | 2485169 | 4.28 |

**Figure 4.**
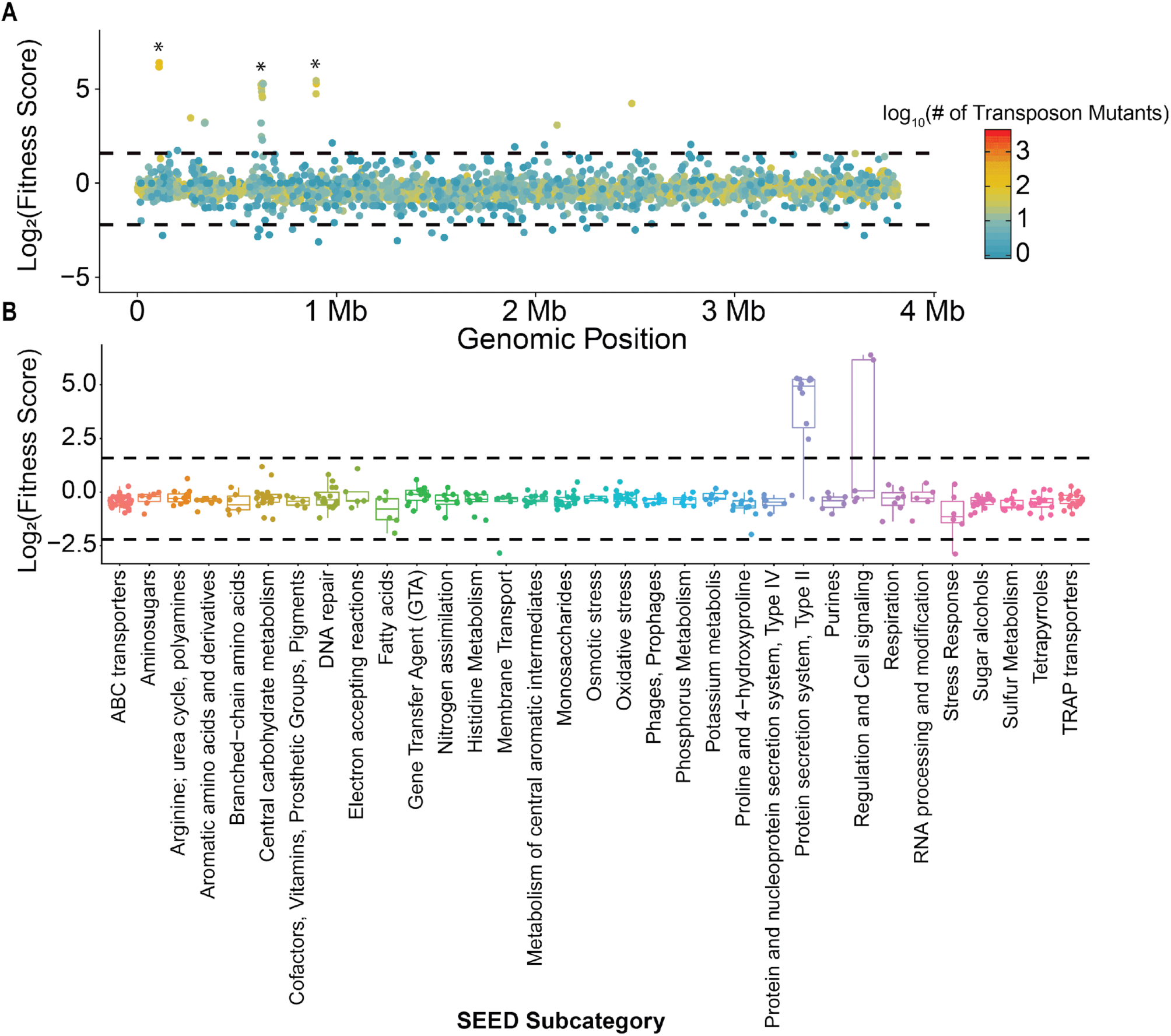
Enriched variants cluster by operon and gene ontology annotation. **A)** Fitness of mutants across the *P. inhibens* genome. Dashed line indicates significance threshold, three standard deviations greater than the mean fitness score. Asterisks indicate location of operons whose mutations exhibited significantly higher fitness scores. Fill color indicates number of unique mutant variants tested for each gene. **B)** SEED subcategory annotations indicate that genes encoding a type II secretion system and cell signaling pathways alter the fitness of *P. inhibens* in the context of phage MD18 infection. Dashed line indicates significance threshold, set at three standard deviations greater than the mean fitness score.

To identify potential mechanistic associations between these significant hits, we used the STRING database to construct a genetic interaction network which identifies associations within lists of genes based on relative genome positions, known experimental studies, coexpression, and cooccurrence in genomes (**Figure 5**)(28). This analysis revealed three primary genetic modules associated with susceptibility to *Phaeobacter* phage MD18. Although most genes were related to the formation of type IV pilus (T4P), hits also included the *chvI*/*chvG* two-component system, and regulators of cellular morphology during cell division.

**Figure 5.**
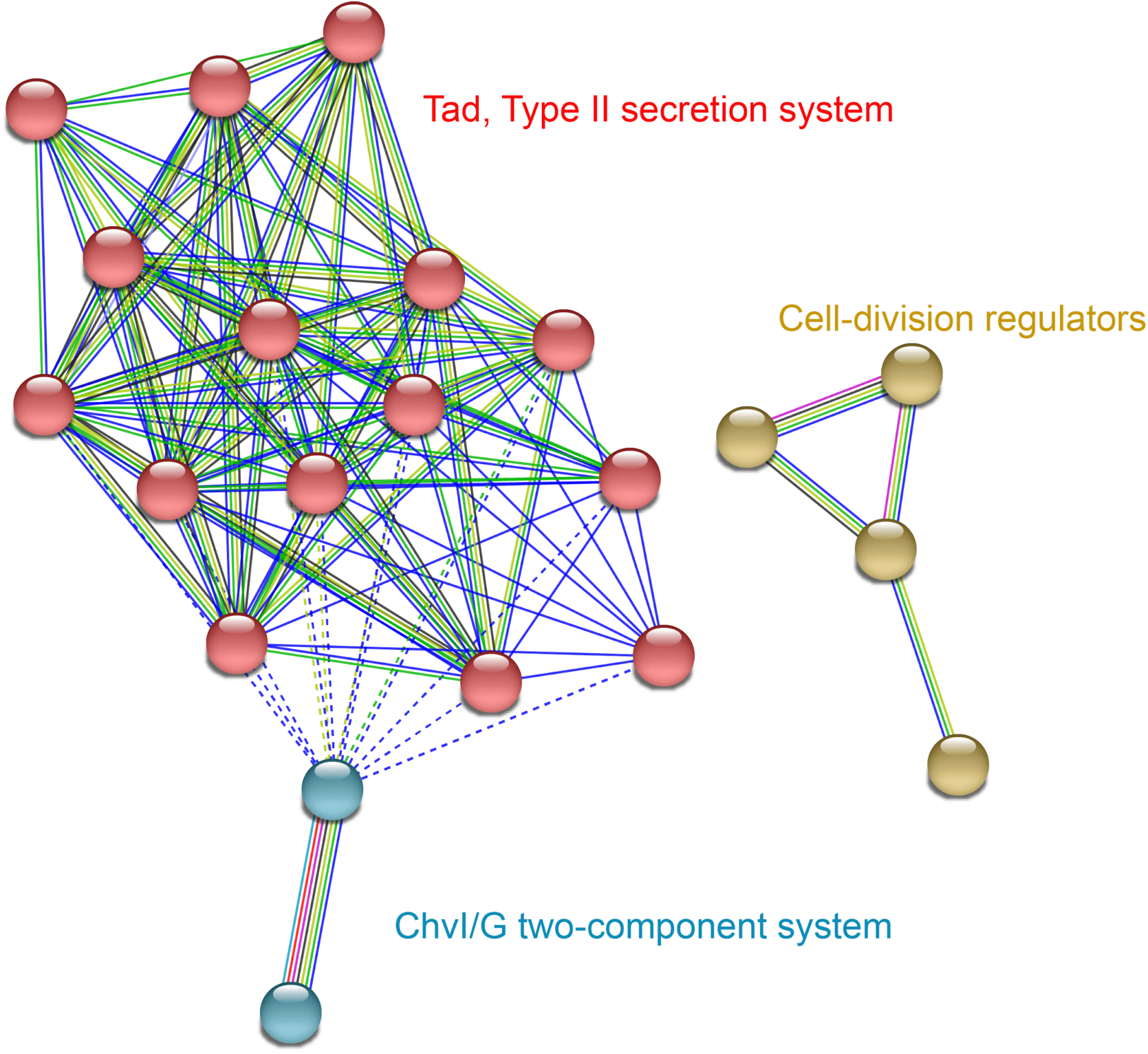
Functional protein association network identifies genetic modules within gene hits. Significant BarSeq hits show enrichment for cellular functions, especially formation of the Tad type II secretion system machinery. Genes were grouped using kmeans clustering, forming three groups (colored). Network construction and visualization were performed using STRING(28). Edges are colored to indicate evidence for interaction and include: interactions known from curated databases (light blue) or experimentally verified (violet); predicted by genomic context such as gene neighborhood (green), gene fusions (red), and gene co-occurence (blue); or predicted based on text mining (yellow-green) or co-expression (black).

## Discussion

In this study, we identified a novel lytic bacteriophage, *Phaeobacter* phage MD18, that infects *P. inhibens*. Based on the genome sequence and morphological characteristics, we propose that MD18 is of the *Siphoviridae* morphotype and constitutes a novel viral genus, which we name ‘*Kasiahvirus*’.

We also identified genetic determinants to MD18 infection in *P. inhibens* using BarSeq. BarSeq has proven to be a promising platform for performing rapid reverse genetics screens to dissect molecular pathways in bacteria(27, 29). In this experiment, large bacterial libraries of transposon insertion mutants containing unique DNA labels are generated and challenged with environmental conditions of interest. Sequencing and determining the relative abundance of these genetic barcodes enables the discovery of genes modulating bacterial fitness under selected conditions. Here, used BarSeq to define the genetic basis for *Phaeobacter inhibens* susceptibility to a novel phage, *Phaeobacter* phage MD18. This analysis showed a clear enrichment of proteins serving a role related to the Tad macromolecular transport system, a Type II secretion system involved in pilus assembly(30), a two-component regulatory system and selected Fts proteins. Thus, our work has revealed several testable hypotheses for how phage MD18 interacts with *P. inhibens*, and how phage-host interactions occur in non-model systems.

First is the hypothesis that *Phaeobacter* phage MD18 recognizes type IV pili as its receptor. These surface structures have been found to be common targets of phages(31–33) and several of their components exhibit significant sequence and structural similarities with bacterial Type II secretion systems(34–36). The relationship between Type IV pili and Type II secretion systems is extensive and it has even been shown that overexpression of certain Type II components results in the formation of pili(30, 37). Future efforts could also utilize adsorption and motility assays to investigate a potential fitness tradeoff for the *P. inhibens* host by losing the Tad pilus.

Second, mutating the *chvI*/*chvG* genes encoding a two-component system was the most potent inhibitor of phage susceptibility, however, it is unclear mechanistically how this two-component system plays a role in phage susceptibility and resistance. Previous studies have shown that mutants of the ChvI-ChvG two-component system lose membrane stability in *Rhizobium leguminosarum*, which could conceivably disrupt the formation of the pilus(38). Alternatively, gene network analysis shows that this two-component system may play a role in regulation of the *pho* operon, and other studies have shown that bacteriophages may hijack this pathway to promote phosphorus uptake in the host, thereby increasing the rate of phage assembly(39). Finally, it is possible that the ChvI-ChvG two-component system directly regulates expression of the Tad pilus.

Several other genes that regulate cell division and assembly of the septum, including the *ftsE*/*ftsX(40)* heterodimer and the peptidoglycan-remodeler *amiC(41)*, were also implicated in altering phage susceptibility. Mutants in these genes exhibit altered cellular morphologies, which may also disrupt or alter pilus formation. Some of these mutations have been shown to lead to a filamentous cell morphology(40), a survival phenotype of many bacteria under stress that this study suggests may help in preventing phage predation. However, it has been shown in other cases that filamentation is associated with increased phage susceptibility(42). It is also possible that misregulated peptidoglycan remodeling physically inhibits injection of phage DNA, a process which is not fully elucidated for tailed phages(43).

The isolation, genome sequencing, and receptor identification of *Phaeobacter* phage MD18 enables future work to understand viral infections of *P. inhibens*. Understanding phage-host interactions in diverse non-model systems like *P. inhibens* will further our understanding of basic phage biology in ways that might increase our abilities to implement new phage technologies and therapies.

## Acknowledgements

We thank the Aretha Fiebig and Sean Crosson for providing the BarSeq transposon mutant library, and the staff at the Marine Biological Laboratory, including Kasia Hammar and Maria Bautista for their assistance with TEM, Fatima Hussain for advice on phage enrichment strategies, and Scott Dawson and Sarah Guest for help in sequencing library preparation. This work was supported by funds from the Marine Biological Lab, DOE (DE-SC0016127), NSF (MCB1822263), HHMI (Grant #5600373), a gift from the Simons Foundation, as well as funds to G.U. (HHMI, UCLA Graduate Division), D.C. (Goldschmidt Foundation and a Arthur Klorfein Scholarship) and D.M.H (Helen Hay Whitney Foundation).

---

**Figure S1.**
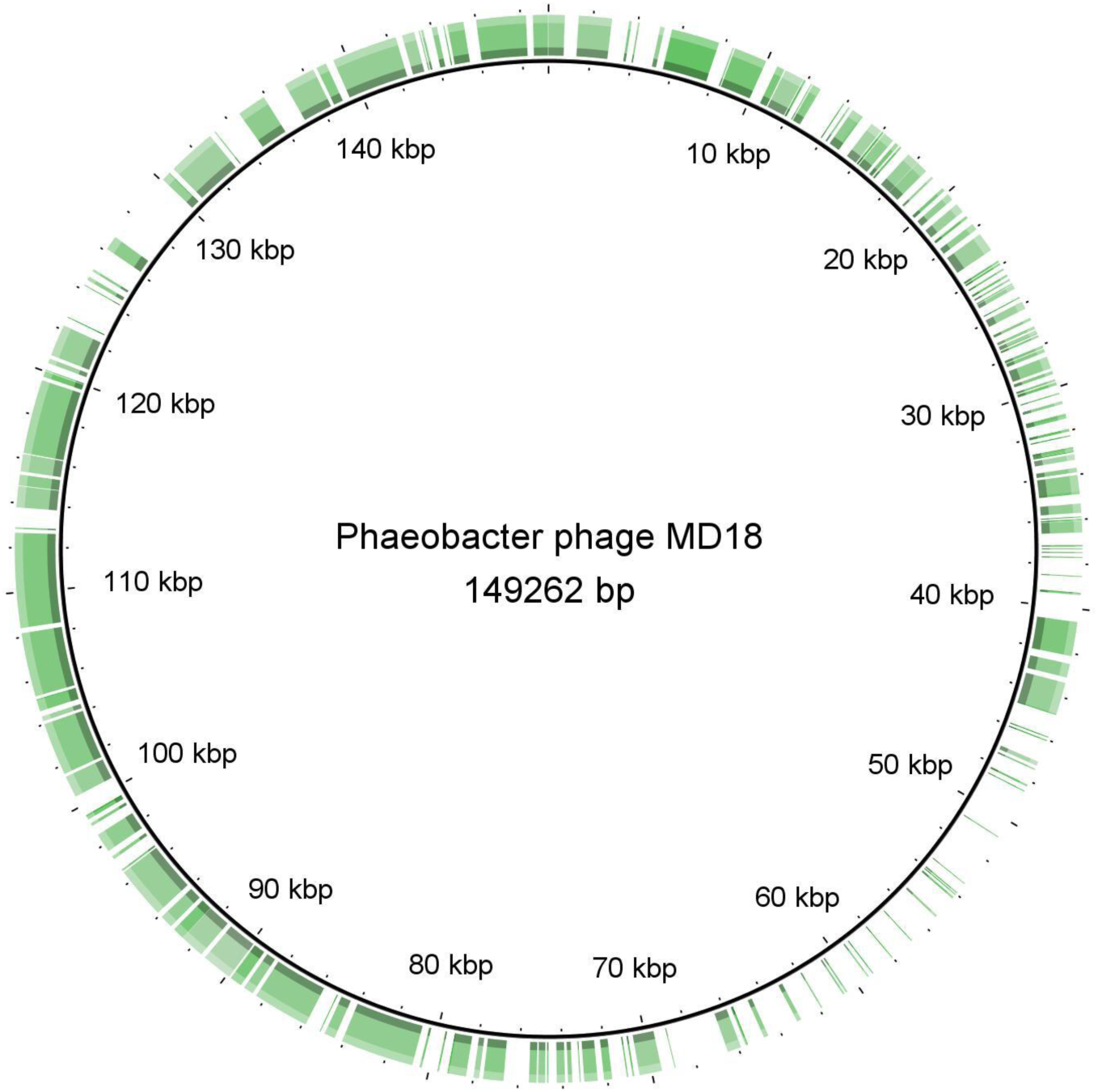
Genomic comparison between *Phaeobacter* phage MD18 genome and *Roseobacter* phage DSS3P8. Sequence alignments of *Phaeobacter* phage MD18 compared to the nearest genomic match, *Roseobacter* phage DSS3P8, visualized using the BLAST Ring Image Generator (BRIG) (44). Highlighted regions indicate sections exhibiting over 50% sequence similarity with *Roseobacter* phage DSS3P8.

**Figure S2.**
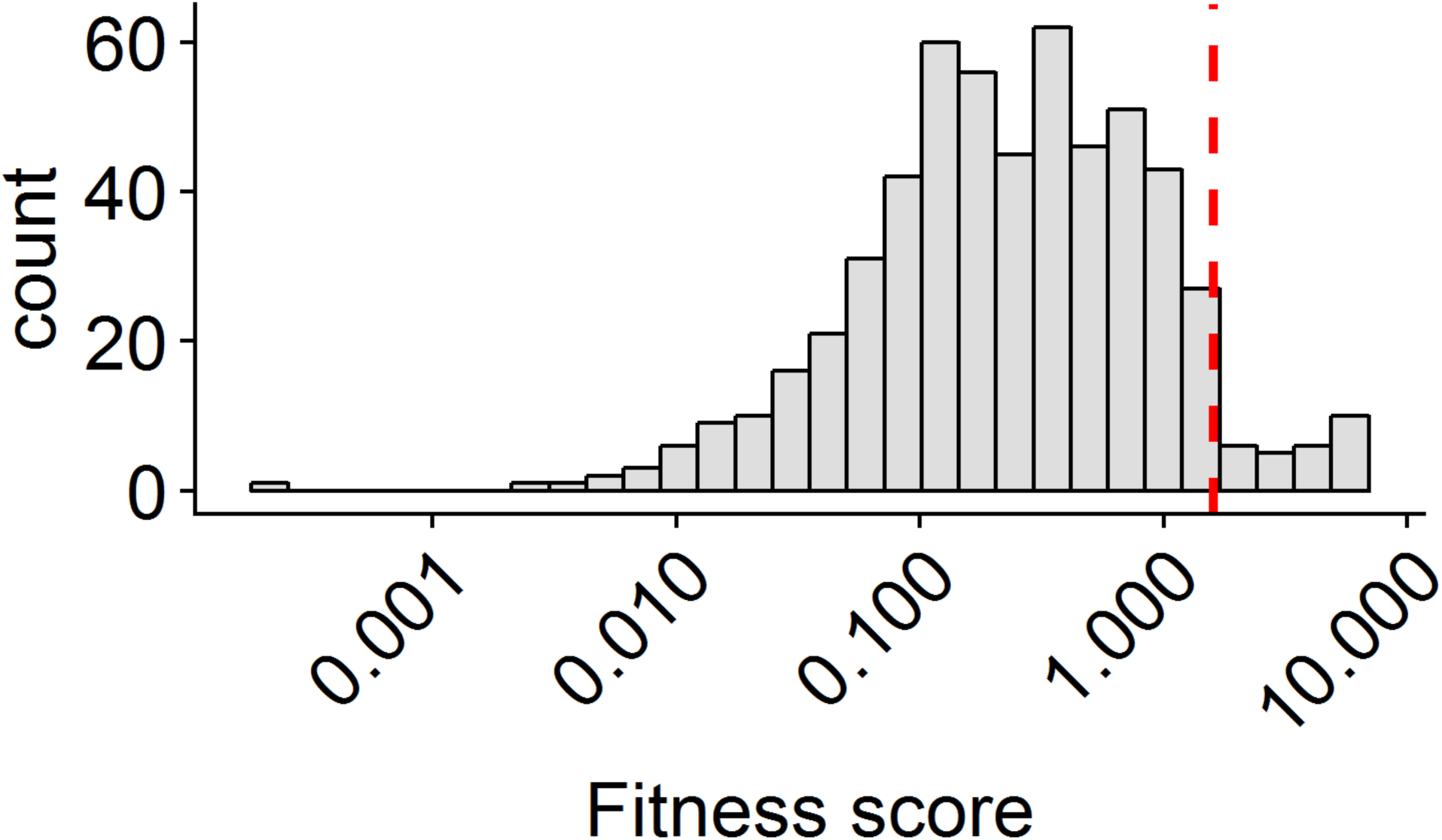
Distribution of gene fitness scores. Gene fitness scores for transposon insertion variants span a wide range. The red line indicates the significance threshold set as three standard deviations greater than the bootstrapped mean of all observed fitness scores.

## Methods

### Bacteriophage isolation and enrichment

*P. inhibens* DSM 17395 was grown in Difco Marine Broth 2216 liquid (MB), on plates supplemented with 1.5% agar, or in 0.5-0.7% top agar overlays at 30°C. The *P. inhibens* infecting bacteriophage was isolated from Woods Hole Waterfront Park in Woods Hole, MA. A sterile 15 mL tube was loaded with a sample of seagrass and filled to 15 mL with seawater. After vigorously vortexing for 5 seconds, 5 mL of this sample was filtered using a 0.22 μm syringe filter and divided into 1 mL aliquots. A single aliquot was incubated overnight at room temperature with 5 mL of log-phase (OD_600_ ∼0.3) *P. inhibens* in MB, centrifuged for 2 minutes at 2,000 x *g*, and filtered using a 0.22 μm syringe filter to obtain the enriched bacteriophage population. The presence of the isolated phage was confirmed by spotting onto 0.7% top agar inoculated with mid-exponential *P. inhibens*. The isolated phage was named “MD18” because it was isolated as part of the Microbial Diversity course at the Marine Biological Laboratory in Woods Hole, MA during the summer of 2018.

### Bacteriophage characterization

Top agar spotting and liquid growth assays were used to measure the concentration of PFUs in the lysate and to characterize the plaque phenotype of the enriched phage MD18 isolate. Eight 10-fold dilutions of the enrichment were prepared by serial dilution in MB. To perform the spotting assay, 4 mL of MB with 0.5% molten agarose was cooled to ∼50°C before mixing with 1 mL of log-phase *P. inhibens*, and this mixture was spread evenly across a petri dish containing MB agar. Once solidified, 5uL of each dilution was spotted and incubated for 16 hrs at 30°C. As a control to exclude the presence of a cellular predators, 10 μL of the undiluted phage stock was incubated with 1 μL 2.0% chloroform for 1 hr with shaking at 4°C before spotting on the same plate. Media control was prepared by incubating 10 μL MB with 1 μL 2.0% chloroform for 1 hr with shaking at 4°C. To perform the liquid growth assays in the presence of bacteriophage, a log-phase culture of *P. inhibens* was diluted 1:100 in MB and 190 μL aliquoted into a 96-well Corning Clear-Bottom plate. To each well, 10 μL of each bacteriophage dilution or sterile MB was added (200 μL total per well) in triplicate. The plate was grown for 24 hrs at 30°C with shaking at 150 rpm and the optical density at 600 nm (OD_600_) of each culture was monitored using a Promega GloMax Explorer Multimode Microplate Reader.

### Transmission electron microscopy

To prepare the bacteriophage for transmission electron microscopy, 10 uL of the enriched bacteriophage isolate lysate (∼10^10^ PFU/ml) was incubated with a glow-discharged formvar coated 200 mesh copper grid for three minutes before being washed three times with sterile 0.22 μm-filtered water. The sample was negatively stained by incubating in a 2% uranyl acetate solution under darkness for 1 minute before undergoing another series of washes. For imaging host cells and phage MD18 together, 5 μL of the same phage preparation was pre-incubated with 5 μL of exponential-phase *P. inhibens* culture for 10 min and adhered to the grid and stained as above. Samples were imaged using a Zeiss 10CA Transmission Electron Microscope with help from the Marine Biological Laboratory Central Microscopy Core.

### Bacteriophage DNA sequencing, assembly, and annotation

Phage MD18 was prepared by inoculating 100 μL of the plaque-isolated phage lysate (∼10^10^ PFU/ml) in 100 mL of mid-log *P. inhibens* in MB. This culture was grown for 24 hr with shaking before centrifugation and 0.22 μm filtration of the supernatant. This phage fraction was concentrated to ∼1 mL with Corning 30,000 MWCO Spin-X UF 20 Concentrators (Corning, NY).

DNA was extracted from this concentrated virus sample using a modified protocol with the Wizard Genomic DNA Purification Kit A1120 (Promega, Madison, WI). Briefly, DNase I was added to 300 uL of phage sample at a final concentration of 1 μg/mL and incubated at room temperature for 1 hr. To the virus sample, 480 μL 50 mM EDTA and 900 μL Cell Lysis Solution were added, vortexed, and incubated for 30 min at 30°C. To this mixture, 600 μL Nuclei Lysis Solution was added, vortexed, and incubated for 5 min at 80°C. To this mixture, 3 μL RNase A Solution was added, vortexed, and incubated for 30 min at 37°C. To this mixture, 200 μL Protein Precipitation Solution was added, vortexed for 20 s, and incubated for 5 min on ice. Cellular debris was pellet by centrifugation at 13,000 x *g* for 3 min, and the pellet discarded. DNA was precipitated by the addition of 2.5 mL isopropanol at -20°C, centrifugation at 13,000 x *g*, and the supernatant discarded. The DNA pellet was washed in 600 μL 95% ethanol at 20°C, centrifuged at 13,000 x *g*, and the supernatant discarded. The DNA pellet was dried before resuspension in 100 μL DNA Rehydration Solution.

*Phaeobacter* phage MD18 DNA was prepared for sequencing with the Nextera DNA Flex Library Prep Kit and barcoded with the IDT for Illumina Nextera DNA UD Indexes (San Deigo, CA). The DNA was sequenced on an Illumina NovaSeq 6000 with a NovaSeq SP reagent kit, yielding 647,478 250-nt read pairs. Reads were trimmed using trimmomatic version 0.38, using the following parameters: SLIDINGWINDOW:4:15 LEADING:2 TRAILING:2 MINLEN:35 resulting in 623,304 reads. These reads were assembled using SPAdes (version 3.11.1) using default parameters. The primary assembled contig was submitted to the RAST(16, 17) online server (https://rast.nmpdr.org/rast.cgi) to identify putative genes and the tRNAscan-SE 2.0(20, 21) online server to identify encoded tRNA genes (http://lowelab.ucsc.edu/tRNAscan-SE/). Genome feature tables were generated by converting Genbank (.gbk) files using GB2sequin online server(45).

### *P. inhibens* barcoded transposon insertion mutant library challenge

The *P. inhibens* library was originally produced by Wetmore *et al*. 2015(27) and generously provided by the Crosson lab. A 1 mL aliquot of this library was used to inoculate 30 mL MB + kanamycin (300ug/uL; kanamycin is the antibiotic resistance marker carried by the transposon used to prepare the BarSeq library) and grown for 4 hours before reaching an OD_600_ of ∼0.5. Three 1.5mL aliquots of this culture were centrifuged at 2,000 x *g* for 2 minutes and the pellets frozen for later DNA preparation. To perform the bacteriophage challenge, 850 uL of the growing culture was used to inoculate six cultures of 30 mL MB + kanamycin (300 μg/μL) and 1.5 mL of 100-fold diluted plaque-purified bacteriophage lysate (∼10^10^ PFU/ml) was added to three of these cultures. These six cultures were grown for eight hours, then 1 mL of each culture was centrifuged at 2,000 x *g* for 2 minutes, and the pellets frozen for later DNA preparation.

### BarSeq sequencing library preparation

In total, nine samples were processed for BarSeq. Three of these samples represented the original library, three represented the library grown without phage, and three more represented the library grown in the presence of phage. Genomic DNA from all samples was harvested using a Promega Maxwell RSC PureFood GMO and Authentication Kit. To prepare these samples for Illumina sequencing, sequencing primer sites, Illumina flow-cell adapters, and sample indices were added using two successive PCRs. Primers used are described in **Table 2**. The first PCR amplifies barcoded transposon sequences from the genomic insertion sites and the second PCR adds Illumina sequencing primer sites and flow cell adapter sequences. For the first PCR, 200 ng of each gDNA sample was amplified using Promega GoTaq G2 Hot Start Polymerase with primers ‘Barseq_P1_PCR1’ and ‘Barseq_P2_PCR1’ (Table 2). The PCR was performed using the following conditions: 95°C for 5 min and 16 cycles of 95°C for 45 s, 59°C for 30 s, and 72°C for 1 min, followed by a final extension for 2 min at 72°C. The DNA from the 50 μL PCR reactions was purified using 60 μL Agencourt AMPure XP beads following manufacturers protocols (Beckman Coulter no. A63880). For the second PCR, 1 μL of each of the nine samples was amplified and indexed by a unique pair of primers ‘Barseq_P1_S51X’ and ‘Barseq_P2_N72X’ with GoTaq G2 Hot Start Polymerase. The same PCR conditions as before were performed for 10 cycles. The DNA from these PCR reactions was purified using 90 uL Agencourt AMPure XP beads (Beckman Coulter A63880) before being pooled in equimolar ratios and sequenced using an Illumina MiSeq Paired-end 250-cycle kit.

**Table 2.**
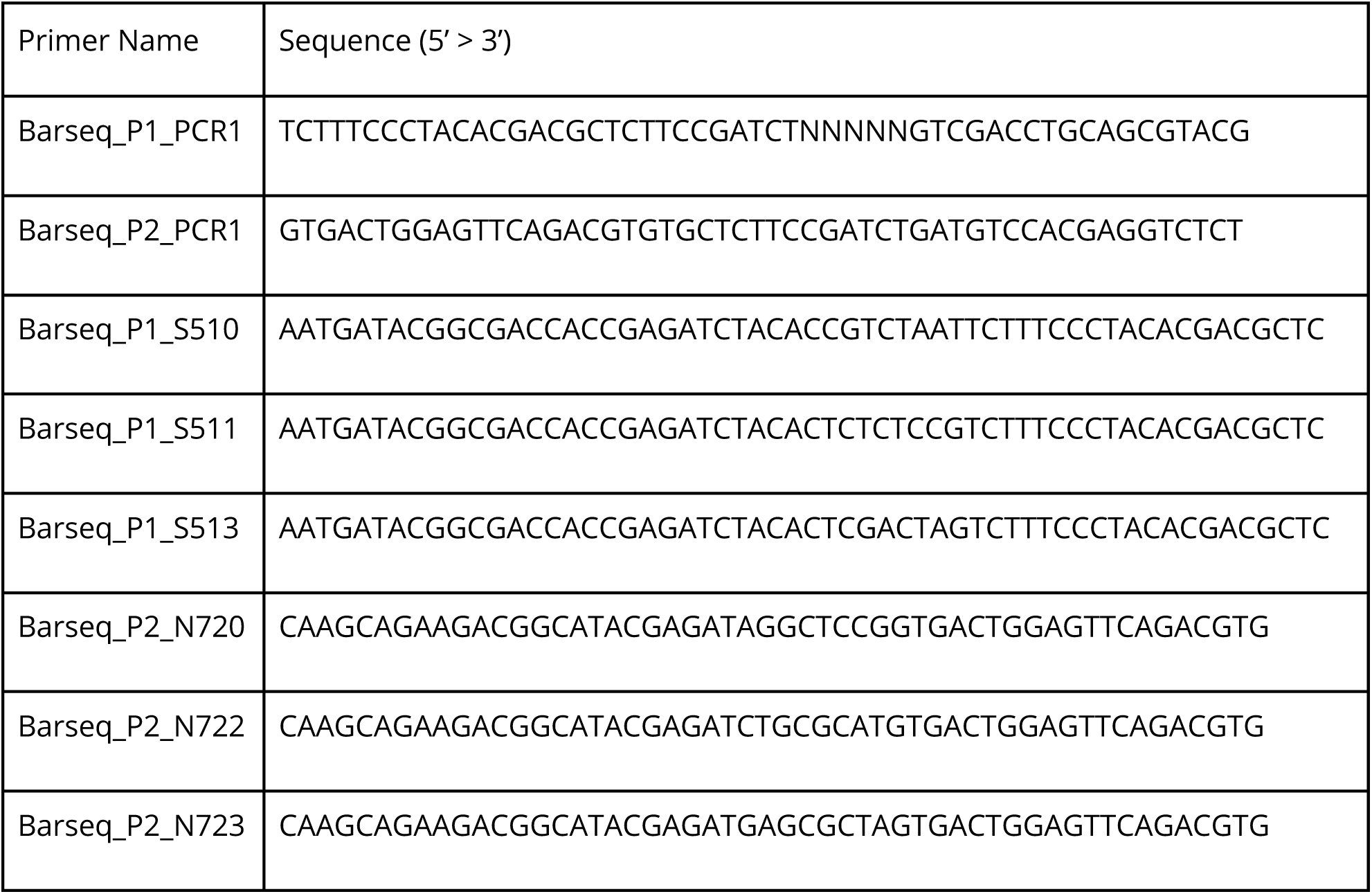
Primers used for BarSeq libraries. Primers used for PCR 1 and PCR 2. PCR 1 added sequencing primers whereas PCR 2 added indices and flow cell adapter sequences.

### Fitness calculation

Transposon insertion mutant fitness was determined by the relative abundance of each barcode before and after challenge by bacteriophage compared to the relative abundance without bacteriophage. Barcode sequences were extracted from the raw reads and the number of each barcode in each sample was counted and normalized to reads per million. The *P. inhibens* DSM 17395 genome (Genbank no. CP002976.1) was annotated using the RAST(16, 17) online server (http://rast.theseed.org/FIG/rast.cgi) and barcodes were grouped by which RAST-identified gene they overlapped. Barcode transposon insertion sites were previously described by Wetmore *et al*. 2015(27). After grouping barcodes by which genes they overlapped, the fitness of each gene was calculated as follows:

First, the gene enrichment score with and without phage were calculated. This was determined by dividing the RPM-normalized counts of all barcode insertions overlapping each gene after treatment by the RPM-normalized counts before treatment.

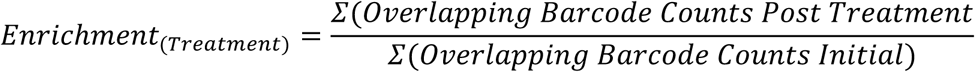

Then, the fitness score of each genetic knockout was calculated by dividing the enrichment score within the phage treatment by their enrichment score without phage added (See equations).

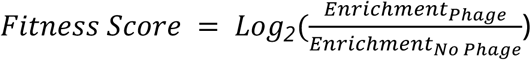

Fitness scores were considered significant if they were greater or less than three standard deviations of the bootstrapped mean fitness score. All gene annotations were generated using RAST with default parameters and viewed using SEED to acquire gene ontology terms. To generate the gene network, amino acid sequences of the genes whose knockouts exhibited significantly greater fitness scores were uploaded as multiple sequences to the STRING online tool (https://string-db.org/) and searched against *Phaeobacter inhibens* DSM 17395. Default settings were followed and online STRING tools were used to identify three gene clusters using K-means clustering.

## Data availability

The *Phaeobacter* phage MD18 genome is available on Genbank (Accession no. MT270409). *P. inhibens* transposon mutant fitness data is available on NCBI Gene Expression Omnibus (Accession no. GSE148502).

